# Differences in IgG antibody responses following BNT162b2 and mRNA-1273 Vaccines

**DOI:** 10.1101/2021.06.18.449086

**Authors:** José G. Montoya, Amy E. Adams, Valérie Bonetti, Sien Deng, Nan A. Link, Suzanne Pertsch, Kjerstie Olson, Martina Li, Ellis C. Dillon, Dominick L. Frosch

**Author notes:** **Corresponding Author:** José G. Montoya M.D., FACP, FIDSA, Dr. Jack S. Remington Laboratory for Specialty Diagnostics, National Reference Laboratory for the Study and Diagnosis of Toxoplasmosis and special pathogens. Palo Alto Medical Foundation, 795 El Camino Real, Ames Building, Palo Alto, CA, USA.

## Abstract

Studies examining antibody responses by vaccine brand are lacking and may be informative for optimizing vaccine selection, dosage, and regimens. The purpose of this study is to assess IgG antibody responses following immunization with BNT162b2 (30 μg S protein) and mRNA-1273 (100 μg S protein) vaccines. A cohort of clinicians at a non-for-profit organization is being assessed clinically and serologically following immunization with BNT162b2 or mRNA-1273. IgG responses were measured at the Remington Laboratory by an IgG against the SARS-CoV-2 spike protein-receptor binding domain. Mixed-effect linear (MEL) regression modeling was used to examine whether the SARS-CoV-2 IgG level differed by vaccine brand, dosage, or days since vaccination. Among 532 SARS-CoV-2 seronegative participants, 530 (99.6%) seroconverted with either vaccine. After adjustments for age and gender MEL regression modeling revealed that the average IgG increased after the second dose compared to the first dose (p<0.001). Overall, titers peaked at week six for both vaccines. Titers were significantly higher for mRNA-1273 vaccine on days 14-20 (p < 0.05), 42-48 (p < 0.01), 70-76 (p < 0.05), 77-83 (p < 0.05), and higher for BNT162b2 vaccine on days 28-34 (p < 0.001). In two participants taking immunosuppressive drugs SARS-CoV-2 IgG remained negative. mRNA-1273 elicited both earlier and higher IgG antibody responses than BNT162b2, possibly due to the higher S-protein delivery. Prospective clinical and serological follow-up of defined cohorts such as this may prove useful in determining antibody protection and whether differences in antibody kinetics between the vaccines have manufacturing relevance and clinical significance.

## INTRODUCTION

Within one year of the emergence of SARS-CoV-2 two novel and effective mRNA vaccines became available, BNT162b2 (Pfizer/BioNTech) and mRNA-1273 (Moderna) (1, 2). BNT162b2 is translated into 30 μg of SARS-CoV-2 full-length spike (pre-fusion conformation) and boosted three weeks after (3). mRNA-1273 is translated into 100 μg of pre-fusion-stabilized spike glycoprotein and boosted four weeks later (4).

Healthcare workers were the first group to receive BNT162b2 and mRNA-1273 (1). The present study was launched on 12/10/20, the week that SARS-CoV-2 vaccines became available, providing the opportunity to assess antibody responses in participants receiving two different vaccine brands, before and after immunization. Most studies so far have focused on following IgG antibody responses to single vaccine brands (5–8). This study examines how antibody responses vary by vaccine brand, dosage, and days since vaccination.

## MATERIALS AND METHODS

A longitudinal study was initiated to estimate the incidence of SARS-CoV-2 infection and COVID-19 by serological testing. Additionally, it aims to assess SARS-CoV-2 antibody responses and sustainability following infection or immunization. Here we report IgG responses following immunization within the first three months of the study. The study protocol was approved by the Sutter Health IRB.

### Serological Assay

Serum SARS-CoV-2 IgG was measured by an automated method (VIDAS^®^ SARS-COV-2 IgG, Biomérieux, France) using an enzyme-linked fluorescent assay (ELFA) at the Dr. Jack S. Remington Laboratory for Specialty Diagnostics at Sutter Health (hereafter Remington Lab, https://www.sutterhealth.org/services/lab-pathology/toxoplasma-serology-laboratory). VIDAS^®^ SARS-COV-2 detects IgG against the receptor binding domain (RBD) of the spike protein. Results are reported as an index (≥ 1.00 = positive). Data from the manufacturer and the Remington lab (n = 199), revealed that this assay had a sensitivity of 100% for specimens obtained ≥ 15 days following onset of symptoms in COVID-19 positive patients. In 989 pre-pandemic samples from the manufacturer, only one tested positive (99.9% specificity)(9).

### Participants

A total of 1,769 clinicians were invited to participate and had to sign the informed consent before enrolling via REDCap. Clinicians belong to a multi-specialty practice comprised of adult and pediatric primary care physicians, specialists (including hospitalists), and advanced practice clinicians. In addition to completing surveys, participants provide serum at baseline and every three months for a year.

### Statistical Analysis

Mixed-effect linear (MEL) regression modeling was used to examine whether the SARS-CoV-2 IgG index measured over time differed by vaccine brand, dosage, or days since vaccination, and examine the interaction effect between the vaccine brand and days since vaccination for the IgG trajectory across time. Modeling adjusted for age and gender and included a subject-specific random intercept term to account for the within-person correlation of measurements over time. The restricted maximum likelihood (REML) approach was used to fit the MEL to produce unbiased estimates of standard errors. Participants who tested positive for SARS-CoV-2 PCR and/or IgG before vaccination (n = 19) were excluded from the model for this report.

## RESULTS

Among 656 clinicians who consented to participate, 611 (93.1%) completed their baseline survey and serum collection. Mean age of participants was 47.4 years. Approximately two-thirds were female (Table 1). Participants self-identified as primarily white (49.8%), Asian (44%), and non-Hispanic (96.2%). Of the 611 participants, 551 (90.2%) completed the three-month follow-up. Of the 551 participants, 532 (96.6%) tested negative for SARS-CoV-2 IgG at baseline and therefore were found eligible for seroconversion. Of the 532 participants, 217 (40.8%) received BNT162b2 and 315 (59.2%) received mRNA-1273.

**Table 1:** Demographics in 611 participants who completed baseline assessment and serum.

| Gender | N | % |
| --- | --- | --- |
| Female | 410 | 67.1% |
| Male | 201 | 32.9% |
| Age mean and categories | 47.4 (9.7) |  |
| 28-39 | 140 | 22.9% |
| 40-49 | 229 | 37.5% |
| 50-59 | 161 | 26.4% |
| 60-76 | 81 | 13.3% |

Seroconversion was demonstrated in 530 (99.6%) of 532 participants. Two participants did not seroconvert following their second dose. In the first non-seroconverting participant, who was receiving a monoclonal antibody (rituximab) against CD20, SARS-CoV-2 antibodies were not detected 28 days following the second dose (BNT162b2) (10). In the second non-seroconverting participant, who was receiving an agent (fingolimod-phosphate) that blocks lymphocytes' ability to emerge from lymph nodes, SARS-CoV-2 antibodies were not detected 21 days following the second dose (mRNA-1273) (11).

Figure 1 depicts the SARS-CoV-2 antibody levels for participants who provided serum samples following vaccination. After adjustments for age and gender, MEL regression modeling found that the IgG increased significantly after the second dose of vaccine compared to the first dose (p<0.001). Overall, titers peaked at week six for both vaccines. Significant differences in IgG were found between vaccine brands, higher for mRNA-1273 on days 14-20 (p < 0.05), 42-48 (p < 0.01), 70-76 (p < 0.05), 77-83 (p < 0.05), and higher for BNT162b2 on days 28-34 (p < 0.001).

**Figure 1. legend.**
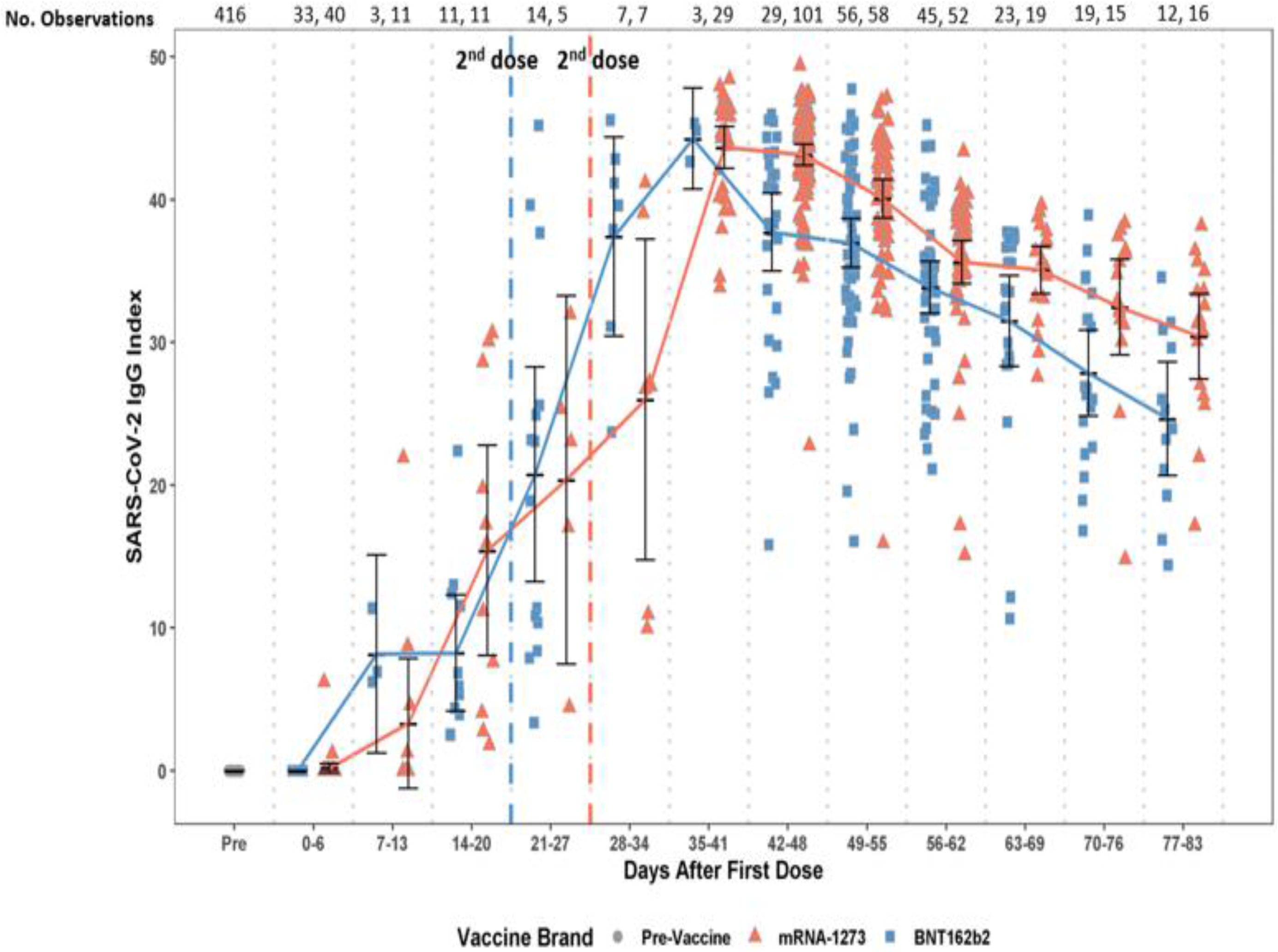
Levels of SARS CoV-2 IgG against the spike protein’s receptor-binding domain by vaccine brand (BNT162b2, mRNA-1273) and over time following first and second doses.

During the days 0-6 post-vaccination, two of 44 participants who received mRNA-1273 had detectable antibodies. In contrast, during the same period, none of the 33 participants who received BNT162b2 had detectable antibodies.

## DISCUSSION

We detected SARS-CoV-2 IgG seroconversion, using an assay aimed at the spike protein-RBD, in all clinicians following either vaccine, with two exceptions who were under immunosuppression. Several differences were identified in the IgG responses to BNT162b2 and mRNA-1273. IgG responses to mRNA-1273 were observed early, within six days following the first dose, while no detectable IgG responses were observed with BNT162b2. IgG responses were more robust to mRNA-1273 than BNT162b2 following the second-dose. It is possible that the higher antigenic load in mRNA-1273, containing more than three-fold the amount of antigen than BNT162b2 explains the significant differences in IgG responses observed. The fact that the antibody kinetics correlated directly with days since vaccination, booster dose, and antigenic content suggests that the mRNA vaccine platforms are suitable for delivery of accurate amounts of antigen despite that it involves translation steps from RNA to protein.

Considering that BNT162b2 is boosted one week earlier than mRNA-1273 may explain why the differences in responses were not even wider in favor of mRNA-1273. If protection requires maintaining antibody levels above a certain threshold, higher initial levels of response following vaccination or frequent boosting may succeed in keeping antibody levels above this threshold for longer than after natural infection (12) despite similar rates of antibody decay. Moreover, even small differences in antibody titers may translate into wider divergence of protection due to amplifiable immune cascades.

Limitations of our paper include that we do not yet have the clinical correlates of immunity that we expect to accrue longitudinally over a minimum of one-year follow-up. Additionally, the clinicians who did not seroconvert due to immunosuppression, did not have measures of T-cell mediated immunity that could still be providing protective immune responses (13). Lastly, this work does not address presence of neutralizing antibodies or antibody responses to other non-mRNA vaccines.

SARS-CoV-2 IgG titers against the spike protein are available to clinical laboratories but have not been studied as surrogate markers for immune protection. Measure of quantitative SARS-CoV-2 spike protein IgG responses plotted over time following immunization in specific cohorts, while tracking clinical correlates, may help to identify individuals who have titer levels that become non-protective. This strategy may serve as the basis to have them studied with other correlates of immune protection (e.g. T-cells) (13) or be candidates for additional doses. To achieve these goals (12, 13), only serological assays targeting the spike protein and with demonstrated sensitivity and specificity, such as the one used for our study, ought to be utilized. Ongoing studies such as ours, can potentially unveil differences in IgG responses between vaccine brands (as observed in this interim report) that may be relevant clinically or for manufacturing purposes (e.g., choice of antigen amount).

## ACKNOWLEDGMENTS

We thank the clinicians who have volunteered and are participating in our study, Palo Alto Medical Foundation/Bay Area Medical Foundation/Sutter Health for their administrative support, and Palo Alto Medical Foundation donors for their financial support.

## REFERENCES

1. Dooling K, Marin M, Wallace M, McClung N, Chamberland M, Lee GM, Talbot HK, Romero JR, Bell BP, Oliver SE. 2021. The Advisory Committee on Immunization Practices' Updated Interim Recommendation for Allocation of COVID-19 Vaccine - United States, December 2020. MMWR Morb Mortal Wkly Rep 69:1657–1660.

2. Rubin EJ, Baden LR, Morrissey S. 2021. Audio Interview: Advice for Clinicians on Covid-19 Vaccines and Social Restrictions. N Engl J Med 384:e77.

3. Polack FP, Thomas SJ, Kitchin N, Absalon J, Gurtman A, Lockhart S, Perez JL, Perez Marc G, Moreira ED, Zerbini C, Bailey R, Swanson KA, Roychoudhury S, Koury K, Li P, Kalina WV, Cooper D, Frenck RW, Jr., Hammitt LL, Tureci O, Nell H, Schaefer A, Unal S, Tresnan DB, Mather S, Dormitzer PR, Sahin U, Jansen KU, Gruber WC, Group CCT. 2020. Safety and Efficacy of the BNT162b2 mRNA Covid-19 Vaccine. N Engl J Med 383:2603–2615.

4. Baden LR, El Sahly HM, Essink B, Kotloff K, Frey S, Novak R, Diemert D, Spector SA, Rouphael N, Creech CB, McGettigan J, Khetan S, Segall N, Solis J, Brosz A, Fierro C, Schwartz H, Neuzil K, Corey L, Gilbert P, Janes H, Follmann D, Marovich M, Mascola J, Polakowski L, Ledgerwood J, Graham BS, Bennett H, Pajon R, Knightly C, Leav B, Deng W, Zhou H, Han S, Ivarsson M, Miller J, Zaks T, Group CS. 2021. Efficacy and Safety of the mRNA-1273 SARS-CoV-2 Vaccine. N Engl J Med 384:403–416.

5. Folegatti PM, Ewer KJ, Aley PK, Angus B, Becker S, Belij-Rammerstorfer S, Bellamy D, Bibi S, Bittaye M, Clutterbuck EA, Dold C, Faust SN, Finn A, Flaxman AL, Hallis B, Heath P, Jenkin D, Lazarus R, Makinson R, Minassian AM, Pollock KM, Ramasamy M, Robinson H, Snape M, Tarrant R, Voysey M, Green C, Douglas AD, Hill AVS, Lambe T, Gilbert SC, Pollard AJ, Oxford CVTG. 2020. Safety and immunogenicity of the ChAdOx1 nCoV-19 vaccine against SARS-CoV-2: a preliminary report of a phase 1/2, single-blind, randomised controlled trial. Lancet 396:467–478.

6. Samanovic MI, Cornelius AR, Wilson JP, Karmacharya T, Gray-Gaillard SL, Allen JR, Hyman SW, Moritz G, Ali M, Koralov SB, Mulligan MJ, Herati RS. 2021. Poor antigen-specific responses to the second BNT162b2 mRNA vaccine dose in SARS-CoV-2-experienced individuals. medRxiv doi:10.1101/2021.02.07.21251311.

7. Stephenson KE, Le Gars M, Sadoff J, de Groot AM, Heerwegh D, Truyers C, Atyeo C, Loos C, Chandrashekar A, McMahan K, Tostanoski LH, Yu J, Gebre MS, Jacob-Dolan C, Li Z, Patel S, Peter L, Liu J, Borducchi EN, Nkolola JP, Souza M, Tan CS, Zash R, Julg B, Nathavitharana RR, Shapiro RL, Azim AA, Alonso CD, Jaegle K, Ansel JL, Kanjilal DG, Guiney CJ, Bradshaw C, Tyler A, Makoni T, Yanosick KE, Seaman MS, Lauffenburger DA, Alter G, Struyf F, Douoguih M, Van Hoof J, Schuitemaker H, Barouch DH. 2021. Immunogenicity of the Ad26.COV2.S Vaccine for COVID-19. JAMA 325:1535–1544.

8. Chu L, McPhee R, Huang W, Bennett H, Pajon R, Nestorova B, Leav B, m RNASG. 2021. A preliminary report of a randomized controlled phase 2 trial of the safety and immunogenicity of mRNA-1273 SARS-CoV-2 vaccine. Vaccine 39:2791–2799.

9. BIOMÉRIEUX. 2021. VIDAS^®^ SARS-COV-2 IgG. https://www.fda.gov/media/140937/download. Accessed

10. Deepak P, Kim W, Paley MA, Yang M, Carvidi AB, El-Qunni AA, Haile A, Huang K, Kinnett B, Liebeskind MJ, Liu Z, McMorrow LE, Paez D, Perantie DC, Schriefer RE, Sides SE, Thapa M, Gergely M, Abushamma S, Klebert M, Mitchell L, Nix D, Graf J, Taylor KE, Chahin S, Ciorba MA, Katz P, Matloubian M, O’Halloran JA, Presti RM, Wu GF, Whelan SPJ, Buchser WJ, Gensler LS, Nakamura MC, Ellebedy AH, Kim AHJ. 2021. Glucocorticoids and B Cell Depleting Agents Substantially Impair Immunogenicity of mRNA Vaccines to SARS-CoV-2. medRxiv doi:10.1101/2021.04.05.21254656.

11. Mehling M, Johnson TA, Antel J, Kappos L, Bar-Or A. 2011. Clinical immunology of the sphingosine 1-phosphate receptor modulator fingolimod (FTY720) in multiple sclerosis. Neurology 76:S20–7.

12. Cromer D, Juno JA, Khoury D, Reynaldi A, Wheatley AK, Kent SJ, Davenport MP. 2021. Prospects for durable immune control of SARS-CoV-2 and prevention of reinfection. Nat Rev Immunol doi:10.1038/s41577-021-00550-x.

13. Sette A, Crotty S. 2021. Adaptive immunity to SARS-CoV-2 and COVID-19. Cell 184:861–880.

